# Shared mechanisms between coronary heart disease and depression: findings from a large UK general population-based cohort

**DOI:** 10.1101/533828

**Authors:** Golam M Khandaker, Verena Zuber, Jessica MB Rees, Livia Carvalho, Amy M Mason, Christopher N Foley, Apostolos Gkatzionis, Peter B Jones, Stephen Burgess

## Abstract

While comorbidity between coronary heart disease (CHD) and depression is evident, it is unclear whether the two diseases have shared underlying mechanisms. We performed a range of analyses in 367,703 unrelated middle-aged participants of European ancestry from UK Biobank, a population based cohort study, to assess whether comorbidity is primarily due to genetic or environmental factors, and to test whether cardiovascular risk factors and CHD are likely to be causally related to depression using Mendelian randomization. We showed family history of heart disease was associated with a 20% increase in depression risk (95% confidence interval [CI] 16% to 24%, p<0.0001), but a genetic risk score that is strongly associated with CHD risk was not associated with depression. An increase of one standard deviation in the CH D genetic risk score was associated with 71% higher CHD risk, but 1% higher depression risk (95% CI 0% to 3%; p=0.11). Mendelian randomization analyses suggested that triglycerides, interleukin-6 (IL-6), and C-reactive protein (CRP) are likely causal risk factors for depression. The odds ratio for depression per standard deviation increase in genetically-predicted triglycerides was 1.18 (95% CI 1.09 to 1.27; p=2×10^-5^); per unit increase in genetically-predicted log-transformed I L-6 was 0.74 (95% CI 0.62 to 0.89; p=0.0012); and per unit increase in genetically-predicted log-transformed CRP was 1.18 (95% CI 1.07 to 1.29; p=0.0009). Our analyses suggest that comorbidity between depression and CHD arises largely from shared environmental factors. I L-6, CRP and triglycerides, are likely to be causally linked with depression, so could be targets for treatment and prevention of depression.

## INTRODUCTION

Coronary heart disease (CHD) and depression are leading causes of disability in high-income countries, and are expected to become so globally by 2030^1,2^. There are three extensively replicated epidemiological observations regarding CHD and depression. First, these conditions are highly comorbid^3^. Second, depression is associated with increased risk of incident CHD^4,5^ and *vice versa^6,7^*. Third, depression is a strong predictor of poor prognosis in people with CHD^4,8^. However, there are also key unanswered questions particularly regarding potential mechanisms underlying this comorbidity. It is unclear whether the association between CHD and depression arises from largely shared genetic or environmental factors. Additionally, while risk factors for CHD are associated with depression in young and older adults^10,11^, it is unclear whether these associations are causal. It is possible that the two illnesses are underpinned by one (or more) shared pathophysiologic mechanism, which manifests as distinct conditions in different organs (i.e., brain and heart).

It was first reported over fifty years ago that around 40% of patients report depression after acute myocardial infarction^12^. Features of mild depression are present in up to two-thirds of patients after acute myocardial infarction^13^, while severe depression is found in around 15% of CHD patients^14^. This association cannot be explained simply as a reaction to emotional trauma from a potentially life-threatening illness. It is clear that the relationship between CHD and depression has bidirectional aspects. Psychological factors including depression were strongly associated with myocardial infarction in the large, multinational, case-control INTERHEART study^15^. Meta-analyses of longitudinal studies confirm that depression is associated with incident CHD after controlling for life-style and other factors^4,5^. Risk factors for CHD such as hyperlipidaemia, hypertension, diabetes and inflammatory markers are associated with risk of depression^9,10^. These findings indicate that CHD and depression may have shared mechanisms.

However, common illnesses such as depression and CHD tend to cluster at population and individual levels^16^, so to what extent this comorbidity is attributable to shared environmental or shared genetic factors is an outstanding question. Residual confounding could be an alternative explanation for previously reported associations between CHD risk factors and depression. Mendelian randomization is an epidemiological approach that uses genetic variants as instrumental variables to untangle the problems of reverse causation (as genetic variants are fixed at conception, hence genetically-predicted levels of risk factors must precede any event) and unmeasured confounding (as genetic variants are often specific in their associations with risk factors).^17^ If genetically-predicted values of a risk factor are associated with a disease outcome, then it is likely the association between the risk factor and outcome has a causal basis.^18^ Discovering common causal risk factors is important clinically as they could inform strategies for primary and secondary prevention. They could also inform further research into shared mechanisms for these comorbid conditions.

We have used UK Biobank, a large population-based cohort study in the United Kingdom, to examine links between probable lifetime major depression (moderate/severe) and risk of CHD. Our investigation consisted of three components. First, we assessed whether family history of heart disease was a predictor of depression. Second, we assessed whether genetic predisposition to CH D risk was a predictor of depression. Third, we performed Mendelian randomization analyses for CHD and various CHD risk factors to determine whether any of these was a causal risk factor for depression.

## SUBJECTS AND METHODS

### Data sources

The UK Biobank cohort comprises around 500,000 participants aged 40 to 69 years at baseline, recruited between 2006-2010 in 22 assessment centres throughout the UK, and followed up for a variety of health conditions from their recruitment date until February 17, 2016 or their date of death^19^. Informed consent was obtained from all participants. The full dataset includes genome-wide genotyping of baseline samples from all participants, results of clinical examinations, assays of biological samples, detailed information on self-reported health behaviour, and is supplemented by linkage with electronic health records such as hospital inpatient data, mortality data and cancer registries. UK Biobank ethical approval is provided by the UK Biobank research ethics committee and Human Tissue Authority research tissue bank. An independent Ethics and Governance Council oversees adherence to the Ethics and Governance Framework and provides advice on the interests of research participants and the general public in relation to UK Biobank. The current study was approved by UK Biobank (ref no. 26999).

We restricted our attention to 367,703 unrelated participants of European ancestry that passed various quality control tests. European ancestry was defined using self-reported ethnicity and genomic principal components as described previously^20^. An initial probable European subset was defined using genomic principal components. Genetic variants were then dropped from the investigation if they had low call rate (3 standard deviations away from the mean) or failed Hardy— Weinberg equilibrium (p-value <10^-6^ for common variants with minor allele frequency >0.01, p-value <10^-12^ for rare variants with minor allele frequency <0.01). Individuals were dropped from the investigation if they had low call rate or excess heterozygosity (3 standard deviations away from the mean), or mismatch between genetic and reported sex. Principal components were then re-calculated on the European ancestry subset, and a further principal component threshold was applied to exclude non-Europeans. Finally, we removed related individuals so that only one person in each family (defined as third-degree relatives or closer) was included in the analysis.

### Outcome

Our primary outcome was self-reported probable lifetime major depression, either moderate or severe, as previously used in UK Biobank^21^.

*Probable moderate lifetime major depression* was defined using four criteria: 1) answering yes to the question “Looking back over your life, have you ever had a time when you were feeling depressed or down for at least a whole week?” or “Have you ever had a time when you were uninterested in things or unable to enjoy the things you used to for at least a whole week?”, 2) answering 2 weeks or more to the question “How many weeks was the longest period when you were feeling depressed or down?”, 3) answering 2 or more to the question “How many periods have you had when you were feeling depressed or down for at least a whole week?”, 4) answering yes to the question “Have you ever seen a general practitioner for nerves, anxiety, tension or depression?”.

*Probable severe lifetime major depression* was defined by the first three criteria and by answering yes to the question “Have you ever seen a psychiatrist for nerves, anxiety, tension or depression?”.

We also considered moderate depression and severe depression separately as secondary outcomes.

### Family history of heart disease and depression

Participants were asked about illnesses of their father and mother, and given a list of conditions to choose from. Family history of heart disease was defined as the participant selecting “heart disease” for either their father or mother. This variable was self-reported, and no validation was attempted.

### Genetic predisposition to coronary heart disease and depression

Genetic risk scores for CH D were calculated using 1.7 million genetic variants and their associations with CHD measured in 60,801 cases and 123,504 controls as described previously^22^. This dataset did not contain any of the UK Biobank participants. This score has been shown to have similar predictive ability to a conventional cardiovascular risk factor score comprising smoking, diabetes, body mass index (BMI), and hypertension (C-index 0.623 for the genetic score versus 0.639 for the risk factor score).

### Mendelian randomization analyses

Mendelian randomization analyses were conducted for i) CHD risk; ii) various conventional cardiovascular risk factors: BMI, waist-hip ratio (WHR), systolic blood pressure (SBP), diastolic blood pressure (DBP), and lipids (low-density lipoprotein [LDL]-cholesterol, high-density lipoprotein [HDL]-cholesterol, and triglycerides); and iii) inflammatory markers: interleukin-1 (IL-1), interleukin-6 (IL-6), fibrinogen, tumour necrosis factor (TNF) alpha, C-reactive protein (CRP), intercellular adhesion molecule 1 (ICAM1), and P-selectin.

### Selection of genetic variants for Mendelian randomization

For CHD risk, we selected 55 genetic variants previously associated with CHD risk at a genome-wide level of significance in a large meta-analysis of the CARDIoGRAMplusC4D consortium^23^ (Supplementary Table 1). Genetic associations with CHD risk were estimated in up to 60,801 CAD cases and 123,504 controls, mostly (∼90%) of European ancestry.

For BMI, we selected 97 genetic variants previously associated with BMI at a genome-wide level of significance in a large meta-analysis of the Genetic Investigation of ANthropometric Traits (GIANT) consortium^24^ (Supplementary Table 2). Genetic associations with BMI are estimated in up to 339,224 participants mostly (95%) of European ancestry. Associations with WHR were obtained for 48 genetic variants associated with WHR at a genome-wide significance level from the same participants in the GIANT consortium^24^ (Supplementary Table 3).

For blood pressure, we selected 93 genetic variants previously associated with either SBP, DBP, or pulse pressure, at a genome-wide level of significance in a large meta-analysis of the UK Biobank CardioMetabolic Consortium BP working group^25^ (Supplementary Table 4). Genetic associations with SBP and DBP are estimated in UK Biobank, in the same 367,703 unrelated participants of European ancestry as the genetic associations with depression.

For lipids, we selected 185 genetic variants previously associated with either LDL-cholesterol, HDL-cholesterol, and triglycerides, at a genome-wide level of significance in a large meta-analysis of the Global Lipids Genetic Consortium^26^ (Supplementary Table 5). Genetic associations with lipids are estimated in up to 188,577 individuals of European ancestry.

For inflammatory markers, we selected genetic variants in the relevant coding gene region previously shown to be conditionally associated with the inflammatory biomarker and only moderately correlated (r^2^<0.6). For interleukin-1, we selected 2 variants (rs6743376 and rs1542176) in the *IL1RN* gene region. For interleukin-6, we selected 3 variants (rs7529229, rs4845371, and rs12740969) in the *IL6R* gene region. For fibrinogen, we selected 1 variant (rs7439150) in the *FGB* gene region. For TNF-alpha, we selected 1 variant (rs1800629) in the *TNF* gene region. For CRP, we selected 4 variants (rs1205, rs3093077, rsll30864, and rsl800947) in the *CRP* gene region. For ICAM1, we selected 5 variants (rs1799969, rs5498, rs1801714, rs281437, and rsll575074) in the *ICAM1* gene region. For P-selectin, we selected 1 variant (rs6136) in the *SELP* gene region. Genetic associations with the biomarkers were obtained from the referenced papers and are listed in Supplementary Table 6.

### Statistical analyses

Associations with the depression outcome (moderate/severe, moderate only, or severe only) were assessed by logistic regression for family history of heart disease and for the CHD genetic risk score. Associations of genetic variants with the outcome were estimated by logistic regression with adjustment for the first 10 principal components of ancestry. Analyses were performed in all participants, and in men and women separately.

For each cardiovascular risk factor and CHD, we performed Mendelian randomization using the inverse-variance weighted method by weighted regression of the genetic associations with the outcome on the genetic associations with the risk factor^27^. We also performed the MR-Egger^28^ and weighted median^29^ methods to assess the robustness of findings.

For lipids, we conducted analyses separately for each lipid fraction including all variants associated with that lipid fraction at a genome-wide level of significance (86 variants for HDL-cholesterol, 76 variants for LDL-cholesterol, 51 variants for triglycerides), as well as for all the lipid fractions in a single analysis model using multivariable Mendelian randomization^30^ with all the 185 genetic variants.

For the inflammatory markers, when there are multiple genetic variants, we report the inverse-variance weighted estimate with adjustment for correlation between the variants^27^. The correlation matrix was estimated using participants from the 1000 Genomes project of European ancestry. This method combines the genetic associations with the outcome into a weighted average association scaled by the genetic association with the biomarker measure. When there is a single genetic variant, we report the per allele genetic association with the outcome.

Two-sided p-values are reported throughout, with correction for multiple testing in the Mendelian randomization analyses using a p-value of 0.05/15 = 0.003, given that 15 risk factors were included in the analysis. Statistical analyses were conducted using snptest version 2.5.2 or R version 3.3.2 (”Sincere Pumpkin Patch”). Code for analyses is available from the authors on request.

## RESULTS

### Participant characteristics

A summary of participant characteristics is presented in Table 1. Out of the 367,703 European unrelated participants, 14,701 (4.0%) had probable lifetime major depression (moderate/severe), of whom 8473 (2.3%) were classed as having moderate depression and 6228 (1.7%) as severe.

**Table 1:** Baseline characteristics of UK Biobank participants

| Characteristic | All-participants | No depression diagnosis | Depression diagnosis |
| --- | --- | --- | --- |
| Sample size, No | 367,703 | 353,002 | 14,701 |
| Male (%) | 45.9 | 46.3 | 36.7 |
| Age-at-recruitment (years) | 57.2 (8.0) | 57.2 (8.0) | 55.7 (7.8) |
| Body mass index (kg/m<sup>2</sup>) | 27.4 (4.8) | 27.3 (4.7) | 27.8 (5.2) |
| Systolic blood pressure (mmHg) | 137.7 (18.6) | 137.8 (18.6) | 134.1 (17.9) |
| Diastolic blood pressure (mmHg) | 82.0 (10.1) | 82.0 (10.1) | 81.0 (10.0) |
| Smoking status (%): |  |  |  |
| - Current | 10.3 | 10.2 | 13.5 |
| - Never | 38.8 | 39 | 34.1 |
| - Ex | 50.8 | 50.7 | 52.4 |
| Alcohol status (%): |  |  |  |
| - Regular (>3 times per week) | 45.4 | 45.6 | 41.9 |
| - Occasional (<3 times per week) | 47.8 | 47.7 | 49.3 |
| - Never/Ex | 6.6 | 6.5 | 8.7 |

Variables are given as percentages or mean (standard deviation).

### Family history of heart disease and depression

There was evidence for an association between family history of heart disease and depression (odds ratio [OR] for moderate/severe depression 1.20, 95% confidence interval [CI]: 1.16 to 1.24; moderate only OR 1.26, 95% CI: 1.21 to 1.32; severe only OR 1.15, 95% CI: 1.12 to 1.18; p < 0.0001 for each outcome). Associations for moderate/severe depression were similar when restricting the analysis to men (OR 1.22, 95% CI: 1.16 to 1.29) and women (OR 1.16, 95% CI: 1.11 to 1.21).

### Genetic predisposition to coronary heart disease and depression

A 1 standard deviation increase in the CHD genetic risk score was associated with a 71% increase in CHD risk.^22^ However, there was only weak evidence for an association between the CHD genetic risk score and depression, and the magnitude of the association was small: OR per 1 standard deviation increase in the genetic risk score for major depression (moderate/severe) 1.01, 95% CI: 1.00 to 1.03, p = 0.11; moderate only OR 1.03, 95% CI: 1.01 to 1.05, p = 0.014; severe only OR 0.99, 95% CI: 0.97 to 1.02, p = 0.66. Associations for moderate/severe depression were almost identical for men (OR 1.01, 95% CI: 0.99 or 1.04) and women (OR 1.01, 95% CI: 0.99,1.04).

### Mendelian randomization analyses

For CHD and the conventional cardiovascular risk factors (Table 2), only triglycerides showed any evidence for a causal effect on depression risk, with an odds ratio of 1.18 (95% CI: 1.09 to 1.27, p = 2×10^-5^) per 1 standard deviation increase in genetically-predicted triglycerides from the inverse-variance weighted method (Figure 1a). Simi ar results were observed in the robust methods (Table 2) and for moderate and severe depression considered separately (Supplementary Tables 7 and 8).

**Table 2:** Mendelian randomization estimates for conventional cardiovascular risk factors on probable lifetime major depression (moderate and severe)

| Risk factor | Method | Odds ratio (95% confidence interval) | p-value |
| --- | --- | --- | --- |
| CHD | IVW | 1.06 (0.99 to 1.13) | 0.11 |
|  | MR-Egger | 1.06 (0.94 to 1.18) | 0.34 |
|  | (intercept) | 1.00 (0.99 to 1.01) | 0.99 |
|  | Weighted median | 1.06 (0.97 to 1.15) | 0.28 |
| BMI | IVW | 1.02 (0.90 to 1.14) | 0.78 |
|  | MR-Egger | 1.26 (0.95 to 1.67) | 0.11 |
|  | (intercept) | 0.99 (0.99 to 1.00) | 0.11 |
|  | Weighted median | 1.02 (0.87 to 1.21) | 0.79 |
| WHR | IVW | 1.00 (0.86 to 1.15) | 0.98 |
|  | MR-Egger | 1.07 (0.59 to 1.92) | 0.83 |
|  | (intercept) | 1.00 (0.98 to 1.01) | 0.82 |
|  | Weighted median | 0.97 (0.78 to 1.19) | 0.75 |
| SBP | IVW | 0.99 (0.95 to 1.04) | 0.73 |
|  | MR-Egger | 0.96 (0.86 to 1.07) | 0.48 |
|  | (intercept) | 1.01 (0.98 to 1.03) | 0.53 |
|  | Weighted median | 0.99 (0.93 to 1.05) | 0.68 |
| DBP | IVW | 1.00 (0.95 to 1.04) | 0.83 |
|  | MR-Egger | 0.98 (0.90 to 1.07) | 0.68 |
|  | (intercept) | 1.00 (0.99 to 1.01) | 0.72 |
|  | Weighted median | 1.01 (0.95 to 1.07) | 0.79 |
| LDL-cholesterol | IVW | 1.01 (0.94 to 1.08) | 0.80 |
|  | MR-Egger | 1.02 (0.91 to 1.14) | 0.74 |
|  | (intercept) | 1.00 (0.99 to 1.01) | 0.82 |
|  | Weighted median | 0.98 (0.90 to 1.08) | 0.73 |
|  | Multivariable | 0.99 (0.93 to 1.05) | 0.70 |
| HDL-cholesterol | IVW | 0.97 (0.91 to 1.03) | 0.31 |
|  | MR-Egger | 0.98 (0.88 to 1.08) | 0.69 |
|  | (intercept) | 1.00 (0.99 to 1.00) | 0.71 |
|  | Weighted median | 1.04 (0.94 to 1.15) | 0.44 |
|  | Multivariable | 1.02 (0.96 to 1.10) | 0.49 |
| Triglycerides | IVW | 1.18 (1.09 to 1.27) | $2 \times 10^{-5}$ |
|  | MR-Egger | 1.16 (1.02 to 1.32) | 0.021 |
|  | (intercept) | 1.00 (0.99 to 1.01) | 0.80 |
|  | Weighted median | 1.18 (1.05 to 1.32) | 0.004 |
|  | Multivariable | 1.17 (1.08 to 1.28) | 0.0003 |

Mendelian randomization estimates are odds ratios for depression (95% confidence intervals) for an increase in the risk factor - per unit increase in the log odds of CHD, per 1 standard deviation increase for BMI and lipid fractions, and per 1 mmHg increase for systolic and diastolic blood pressure.

**Figure 1:**
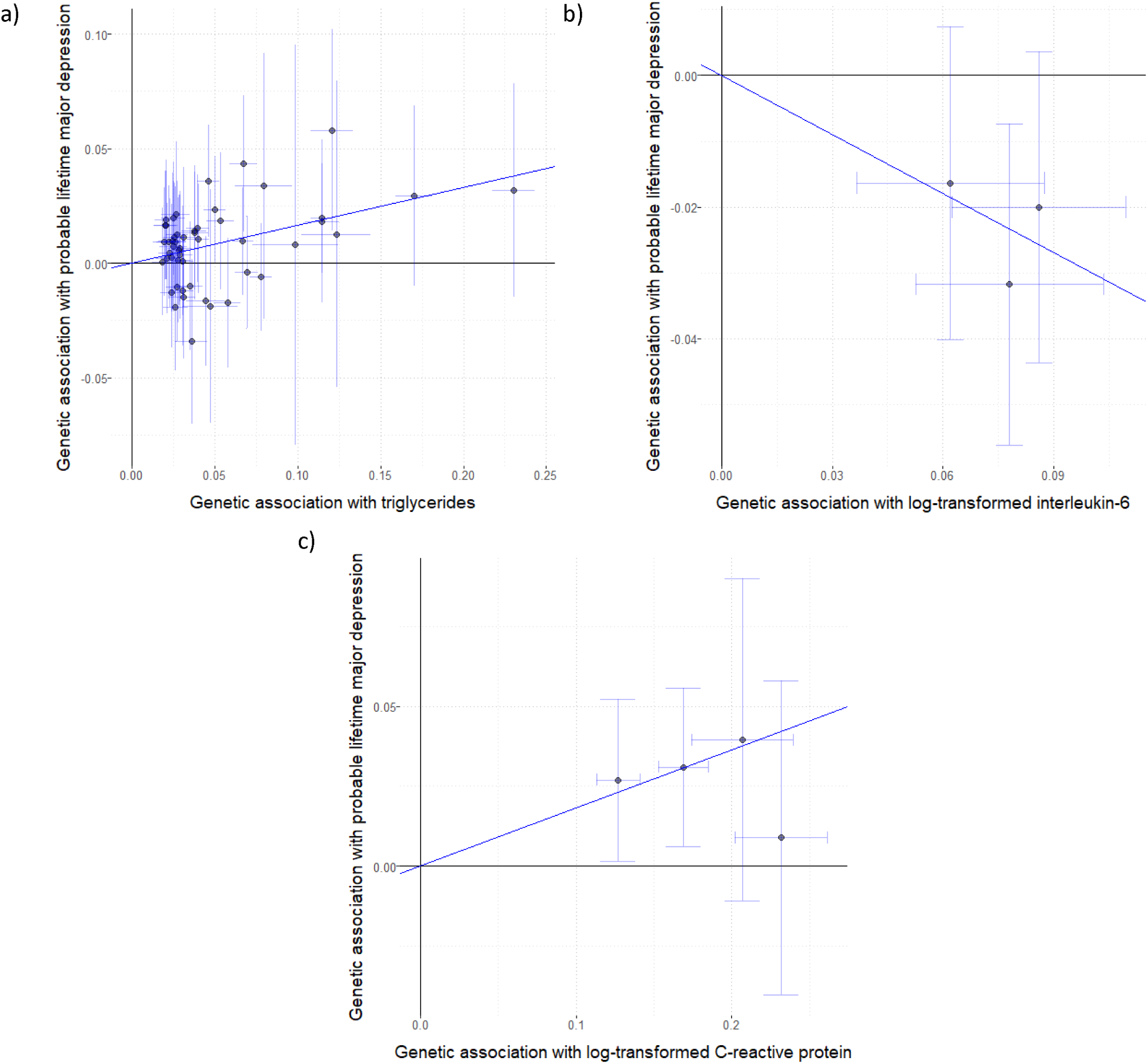
Per allele genetic associations with exposure plotted against associations with risk of probable lifetime major depression, moderate/severe (log odds ratios) for three exposures: a) triglycerides (SD units) based on 51 genome-wide significant predictors; b) interleukin-6 (log-transformed) based on 3 genetic variants located in the *IL6R* gene region; c) C-reactive protein (log-transformed) based on 4 genetic variants located in the *CRP* gene region. The solid line represents the estimate from the inverse-variance weighted method.

For the inflammatory biomarkers (Table 3), there was evidence for causal effects of IL-6 and CRP, with an odds ratio of 0.74 (95% CI: 0.62 to 0.89, p = 0.0012) per unit increase in genetically-predicted values of log-transformed IL-6 (Figure 1b), and 1.18 (95% CI: 1.07 to 1.29, p = 0.0009) per unit increase in genetically-predicted values of log-transformed CRP (Figure 1c). Again, similar results were observed for moderate and severe depression considered separately (Supplementary Tables 9 and 10). All other results were compatible with the null, although there was nominal significance in the association of increased genetically-predicted IL-1 with increased risk of severe depression (p = 0.037).

**Table 3:** Mendelian randomization results for inflammatory biomarkers on probable lifetime major depression (moderate and severe)

| Biomarker | Scale | OR (95% CI) | p-value |
| --- | --- | --- | --- |
| IL-1 | Per 1 SD increase in IL-1Ra | 1.06 (0.98 to 1.14) | 0.17 |
| IL-6 | Per unit increase in log-transformed IL-6 | 0.74 (0.62 to 0.89) | 0.0012 |
| Fibrinogen | Per unit increase in log-transformed fibrinogen | 0.96 (0.42 to 2.20) | 0.92 |
| TNF-alpha | Per TFN expression increasing allele | 0.99 (0.96 to 1.02) | 0.68 |
| CRP | Per 1 unit increase in log-transformed CRP | 1.18 (1.07 to 1.29) | 0.0009 |
| ICAM-1 | Per 20ng/dL increase in sICAM-1 | 1.01 (0.99 to 1.04) | 0.22 |
| P-selectin | Per P-selectin concentration increasing allele | 0.97 (0.95 to 1.00) | 0.16 |

Mendelian randomization estimates are odds ratios for depression (95% confidence intervals) either per unit increase in the risk factor or per allele as indicated.

## DISCUSSION

We performed several analyses to understand potential shared mechanisms between CH D and depression. Our analyses suggest family history of heart disease is strongly associated with probable lifetime major depression (moderate/severe) with a 20% relative increase in depression risk associated with reporting at least one parent dying of heart disease, but a genetic risk score that predicts CHD risk almost as well as conventional risk factors was not strongly associated with depression. This suggests that the comorbidity between the two conditions arises largely from shared environmental factors. Further investigation into cardiovascular risk factors using the Mendelian randomization paradigm elucidated potential biological pathways contributing to comorbidity. We provide evidence that, out of all cardiovascular risk factors, triglycerides and the inflammatory markers IL-6 and CRP are likely to be causally related to depression.

Accumulating evidence supports an important role for low-grade systemic inflammation in depression^31^. Meta-analyses of cross-sectional studies confirm that concentrations of inflammatory markers, such as CRP, IL-6, TNF-alpha, IL-1β, are elevated in peripheral blood during an acute depressive episode^32,33^, then tend to subside after recovery but continue to be elevated in treatment resistant patients^32,34,35^. Population-based longitudinal studies including our own work have reported that higher concentrations of CRP or IL-6 in childhood or adult life are associated with increased risk of depression or persistent depressive symptoms subsequently at follow-up^9,11,36^”^38^. Furthermore, a genetic variant in the *IL6R* gene (Asp358Ala; rs2228145 A>C), which is known to dampen down inflammation by impairing IL-6R signalling^39^, was previously reported to be protective for severe depression^40^. Depressive symptoms and IL-6 share common genetic predictors^41^. Findings from our Mendelian randomization analyses add to the existing literature by showing that reverse causality or residual confounding are unlikely explanations for previously reported associations between IL-6, CRP and depression. We provide evidence that inflammation, particularly the CRP and IL-6/IL-6R pathways, is causally involved in pathogenesis of depression.

A previous MR study did not find evidence for a causal association between depression and CRP^42^. Statistical power could be a reason for the null findings (1128 cases of depression compared with 14,701 in the current analysis). Previous Mendelian randomization investigations have suggested that IL-6 is a causal determinant of CHD risk^43,44^, but this is not the case for CRP^45,46^. Therefore, with regards to shared inflammatory mechanisms underpinning the comorbidity between depression and CHD it is likely that IL-6 is a key driver.

Consistent with our findings, a recent Mendelian randomization study reported potential causal relationships with depression for LDL-cholesterol and for triglycerides^47^. The finding is unlikely to be driven by central obesity, because BMI or WHR are not causally linked with depression. Systematic reviews of observational studies suggest that depression is associated with lower total cholesterol^48^, but the evidence for LDL-cholesterol is mixed^49^. A systematic review of triglyceride concentration in depression compared with controls is currently lacking. Inflammation leads to changes in lipid metabolism including decreased HDL cholesterol and increased triglycerides levels^50^. Anti-inflammatory treatment that inhibits IL-6 also inhibits triglycerides^51^. Elevated IL-6^9^ and triglycerides^52^ levels in childhood are associated with increased risk of depression in young adulthood. However, lowering of serum cholesterol in middle-aged subjects by diet, drugs, or both, leads to a decrease in CHD but an increase in deaths due to suicide or violence^53^. Mechanisms for this effect are not understood, but it has been proposed that low membrane cholesterol could affect serotonergic neurotransmission by decreasing the number of serotonin receptors^53^. Further work is needed to understand the relationships between lipid alterations and depression.

How inflammation causes depression and CHD has been studied extensively; see reviews^54,55^. In brief, direct effects of peripheral inflammation relevant for cardiovascular risk include the development of atherosclerotic lesions in the arterial tree, and effects on endothelial reactivity and myocardial function^55^. Peripheral inflammation also influences the central nervous system. Brain responses to inflammation involve neural systems for motivational and homeostatic control and are expressed through depressed mood state and changes in autonomic cardiovascular regulation^56^. As for depression, extensive animal and human studies have demonstrated that peripheral immune activation leads to changes in mood and behaviour by increasing turnover of serotonin, oxidative stress, activation of the hypothalamic-pituitary-adrenal (HPA) axis, and by reducing synaptic plasticity^54^.

A strong association of depression with family history of heart disease, but not with a genetic risk score for CH D, suggests that the comorbidity between depression and CVD arises largely from shared environmental factors. Although we did not observe an association between the genetic risk score for CHD and depression, it is likely that there is some contribution from shared genetic predictors, as a genetic correlation between CHD and depression has previously been reported. However, the magnitude of this correlation was low^57,58^. According to a large Swedish twin study, acute environmental factors play a large role in depression-CHD comorbidity in men, whereas in women chronic factors, which are in part genetic, are more important^58^. However, we did not observe any sex-difference in the observational associations of depression with CHD family history or genetic risk. Nearly two thirds of depression cases in our study were female. The observational association between family history of CHD and depression may also be influenced by dynastic effects. For example, individuals with family history of heart disease will have been exposed to risk factors for heart disease (and potentially also for depression) through their family environment, i.e. confounding by shared environment.

With regards to shared environmental factors, the depression-CHD comorbidity could be linked with early-life factors influencing inflammatory regulation, such as impaired fetal development or childhood maltreatment. This idea is consistent with the common-cause or fetal programming hypothesis by David Barker^59^. Low birth weight, a catchall marker of suboptimal fetal development, and childhood maltreatment are associated with increased levels of circulating inflammatory markers^60,61^, depression^62^ and CHD^63^ in adulthood.

Limitations of the work include the quality of the depression outcome. A working group of the UK Biobank study used self-reported data to create variables for probable recurrent major depression (moderate/severe) named according to Diagnostic and Statistical Manual for Mental Disorders (DSM) terminology. We use the term probable lifetime major depression to describe the same variable as we could not be certain whether criteria for ‘recurrent’ were fulfilled. Nevertheless, based on these criteria the prevalence of probable lifetime major depression (moderate/severe) in our sample was 4.0%, which is unlikely to be an overestimate. Lifetime prevalence of major depression is around 10-20% according to general population studies^64^. Misclassification of cases as controls introduces a bias towards the null, so the associations between genetic variants and depression observed in our analysis are likely to be conservative, and may well under-estimate the true impact of the risk factors on disease risk. Our investigation focused on CHD and cardiovascular risk factors as causes of depression, but the reverse analysis could also be performed. As future work, the association between a genetic risk score for depression and CHD risk could be investigated.

A better understanding of causal risk factors for depression could inform novel strategies for treatment and prevention. Lifestyle modifications targeting ‘inflammogenic’ risk factors such as physical inactivity, obesity, smoking and alcohol use could improve risks for both CHD and depression. Inflammation is associated with antidepressant resistance^35,65^, so anti-inflammatory drugs may be helpful for patients with depression who show evidence of immune activation. Anti-cytokine drugs that inhibit IL-6 or TNF-alpha signalling improve depressive symptoms in patients with chronic inflammatory illness, independently of improvement in physical illness^66^. Randomized trials testing the efficacy of novel anti-inflammatory drugs in patients with depression are currently ongoing.

In summary, we provide evidence that the comorbidity between depression and CHD largely arises from shared environmental factors. Certain risk factors for CHD, specifically IL-6, CRP and triglycerides, are likely to be causally linked with depression, so these could be targets for treatment and prevention of the illness.

## Acknowledgements

Golam Khandaker acknowledges funding support from the MQ: Transforming Mental Health (Data Science Award; grant code: MQDS17/40), the Wellcome Trust (Intermediate Clinical Fellowship; grant code: 201486/Z/16/Z), and the Medical Research Council (MICA: Mental Health Data Pathfinder; grant code: MC_PC_17213). Stephen Burgess is supported by a Sir Henry Dale Fellowship jointly funded by the Wellcome Trust and the Royal Society (Grant Number 204623/Z/16/Z). The authors thank Michael Inouye for help with data acquisition. This research has been conducted using the UK Biobank Resource under Application Number 26999.

