## Supplementary Material for "Shared mechanisms between coronary heart disease and depression: findings from a large UK general population-based cohort"

Supplementary Table 1: Genetic predictors of CHD included in Mendelian randomization analysis

| rsid | chr:position | rsid | chr:position |
| --- | --- | --- | --- |
| rs7528419 | chr1:109817192 | rs2519093 | chr9:136141870 |
| rs6689306 | chr1:154395946 | rs2891168 | chr9:22098619 |
| rs67180937 | chr1:222823743 | rs11191416 | chr10:104604916 |
| rs11206510 | chr1:55496039 | rs2487928 | chr10:30323892 |
| rs9970807 | chr1:56965664 | rs1870634 | chr10:44480811 |
| rs17678683 | chr2:145286559 | rs1412444 | chr10:91002927 |
| rs16986953 | chr2:19942473 | rs2128739 | chr11:103673277 |
| rs6725887 | chr2:203745885 | rs964184 | chr11:116648917 |
| rs515135 | chr2:21286057 | rs10840293 | chr11:9751196 |
| rs6544713 | chr2:44073881 | rs3184504 | chr12:111884608 |
| rs7568458 | chr2:85788175 | rs2681472 | chr12:90008959 |
| rs9818870 | chr3:138122122 | rs11838776 | chr13:111040681 |
| rs4593108 | chr4:148281001 | rs9319428 | chr13:28973621 |
| rs72689147 | chr4:156639888 | rs10139550 | chr14:100145710 |
| rs17087335 | chr4:57838583 | rs56062135 | chr15:67455630 |
| rs273909 | chr5:131667353 | rs4468572 | chr15:79124475 |
| rs9349379 | chr6:12903957 | rs8042271 | chr15:89574218 |
| rs12202017 | chr6:134173151 | rs17514846 | chr15:91416550 |
| rs55730499 | chr6:161005610 | rs12936587 | chr17:17543722 |
| rs4252185 | chr6:161123451 | rs216172 | chr17:2126504 |
| rs17609940 | chr6:35034800 | rs46522 | chr17:46988597 |
| rs56336142 | chr6:39134099 | rs7212798 | chr17:59013488 |
| rs10953541 | chr7:107244545 | rs663129 | chr18:57838401 |
| rs11556924 | chr7:129663496 | rs56289821 | chr19:11188247 |
| rs3918226 | chr7:150690176 | rs4420638 | chr19:45422946 |
| rs2107595 | chr7:19049388 | rs28451064 | chr21:35593827 |
| rs2954029 | chr8:126490972 | rs180803 | chr22:24658858 |
| rs264 | chr8:19813180 |  |  |

Supplementary Table 2: Genetic predictors of BMI included in Mendelian randomization analysis

| rsid | chr:position | rsid | chr:position |
| --- | --- | --- | --- |
| rs1558902 | chr16:53803574 | rs2650492 | chr16:28333411 |
| rs6567160 | chr18:57829135 | rs6804842 | chr3:25106437 |
| rs13021737 | chr2:632348 | rs12940622 | chr17:78615571 |
| rs10938397 | chr4:45182527 | rs7164727 | chr15:73093991 |
| rs543874 | chr1:177889480 | rs11847697 | chr14:30515112 |
| rs2207139 | chr6:50845490 | rs4740619 | chr9:15634326 |
| rs11030104 | chr11:27684517 | rs492400 | chr2:219349752 |
| rs3101336 | chr1:72751185 | rs13191362 | chr6:163033350 |
| rs7138803 | chr12:50247468 | rs3736485 | chr15:51748610 |
| rs10182181 | chr2:25150296 | rs17001654 | chr4:77129568 |
| rs3888190 | chr16:28889486 | rs11191560 | chr10:104869038 |
| rs1516725 | chr3:185824004 | rs2080454 | chr16:49062590 |
| rs12446632 | chr16:19935389 | rs7715256 | chr5:153537893 |
| rs2287019 | chr19:46202172 | rs2176040 | chr2:227092802 |
| rs16951275 | chr15:68077168 | rs1528435 | chr2:181550962 |
| rs3817334 | chr11:47650993 | rs2075650 | chr19:45395619 |
| rs2112347 | chr5:75015242 | rs1000940 | chr17:5283252 |
| rs12566985 | chr1:75002193 | rs2033529 | chr6:40348653 |
| rs3810291 | chr19:47569003 | rs11583200 | chr1:50559820 |
| rs7141420 | chr14:79899454 | rs7239883 | chr18:40147671 |
| rs13078960 | chr3:85807590 | rs2836754 | chr21:40291740 |
| rs10968576 | chr9:28414339 | rs9400239 | chr6:108977663 |
| rs17024393 | chr1:110154688 | rs10733682 | chr9:129460914 |
| rs657452 | chr1:49589847 | rs11688816 | chr2:63053048 |
| rs12429545 | chr13:54102206 | rs11057405 | chr12:122781897 |
| rs12286929 | chr11:115022404 | rs9914578 | chr17:2005136 |
| rs13107325 | chr4:103188709 | rs977747 | chr1:47684677 |
| rs11165643 | chr1:96924097 | rs2121279 | chr2:143043285 |
| rs7903146 | chr10:114758349 | rs29941 | chr19:34309532 |
| rs10132280 | chr14:25928179 | rs11727676 | chr4:145659064 |
| rs17405819 | chr8:76806584 | rs3849570 | chr3:81792112 |
| rs6091540 | chr20:51087862 | rs9374842 | chr6:120185665 |
| rs1016287 | chr2:59305625 | rs6477694 | chr9:111932342 |
| rs4256980 | chr11:8673939 | rs4787491 | chr16:30015337 |
| rs17094222 | chr10:102395440 | rs1441264 | chr13:79580919 |
| rs12401738 | chr1:78446761 | rs7899106 | chr10:87410904 |
| rs7599312 | chr2:213413231 | rs2176598 | chr11:43864278 |
| rs2365389 | chr3:61236462 | rs2245368 | chr7:76608143 |
| rs205262 | chr6:34563164 | rs17203016 | chr2:208255518 |
| rs2820292 | chr1:201784287 | rs17724992 | chr19:18454825 |
| rs12885454 | chr14:29736838 | rs7243357 | chr18:56883319 |
| rs9641123 | chr7:93197732 | rs16907751 | chr8:81375457 |
| rs12016871 | chr13:28017782 | rs1808579 | chr18:21104888 |
| rs16851483 | chr3:141275436 | rs13201877 | chr6:137675541 |
| rs1167827 | chr7:75163169 | rs2033732 | chr8:85079709 |
| rs758747 | chr16:3627358 | rs9540493 | chr13:66205704 |
| rs1928295 | chr9:120378483 | rs1460676 | chr2:164567689 |
| rs9925964 | chr16:31129895 | rs6465468 | chr7:95169514 |
| rs11126666 | chr2:26928811 |  |  |

Supplementary Table 3: Genetic predictors of WHR included in Mendelian randomization analysis

| rsid | chr:position | rsid | chr:position |
| --- | --- | --- | --- |
| rs2645294 | chr1:119574587 | rs1936805 | chr6:127452116 |
| rs905938 | chr1:154991389 | rs10245353 | chr7:25858614 |
| rs10919388 | chr1:170372503 | rs7801581 | chr7:27223771 |
| rs714515 | chr1:172352990 | rs7830933 | chr8:23603324 |
| rs2820443 | chr1:219753509 | rs12679556 | chr8:72514228 |
| rs1385167 | chr2:66200648 | rs10991437 | chr9:107735920 |
| rs10195252 | chr2:165513091 | rs7917772 | chr10:104487443 |
| rs1569135 | chr2:188115398 | rs11231693 | chr11:63862612 |
| rs17819328 | chr3:12489342 | rs10842707 | chr12:26471364 |
| rs2276824 | chr3:52637486 | rs1443512 | chr12:54342684 |
| rs2371767 | chr3:64718258 | rs4765219 | chr12:124440110 |
| rs10804591 | chr3:129334233 | rs8042543 | chr15:31708263 |
| rs17451107 | chr3:156797609 | rs8030605 | chr15:56504598 |
| rs3805389 | chr4:56482750 | rs1440372 | chr15:67033151 |
| rs9991328 | chr4:89713121 | rs2925979 | chr16:81534790 |
| rs303084 | chr4:124066948 | rs4646404 | chr17:17420199 |
| rs9687846 | chr5:55861894 | rs8066985 | chr17:68453345 |
| rs1045241 | chr5:118729286 | rs12454712 | chr18:60845884 |
| rs7705502 | chr5:173320815 | rs12608504 | chr19:18389135 |
| rs6556301 | chr5:176527577 | rs4081724 | chr19:33824946 |
| rs1294410 | chr6:6738752 | rs979012 | chr20:6623374 |
| rs7759742 | chr6:32381736 | rs224333 | chr20:34023962 |
| rs1358980 | chr6:43764551 |  |  |

Supplementary Table 4: Genetic predictors of blood pressure included in MR analysis

| rsid | chr:position | rsid | chr:position |
| --- | --- | --- | --- |
| rs139385870 | chr1:1685921 | rs67330701 | chr11:69079707 |
| rs3820068 | chr1:15798197 | rs7178615 | chr15:66869072 |
| rs10922502 | chr1:89360158 | rs62012628 | chr15:79070000 |
| rs7562 | chr2:28635740 | rs12906962 | chr15:95312071 |
| rs13420463 | chr2:37517566 | rs12921187 | chr16:4943019 |
| rs55780018 | chr2:208526140 | rs72799341 | chr16:30936743 |
| rs9859176 | chr3:134000025 | rs8059962 | chr16:81574197 |
| rs13112725 | chr4:106911742 | rs4308 | chr17:61559625 |
| rs10059921 | chr5:87514515 | rs745821 | chr18:48142854 |
| rs6595838 | chr5:127868199 | rs62104477 | chr19:30294991 |
| rs6911827 | chr6:22130601 | rs6108168 | chr20:8626271 |
| rs78648104 | chr6:50683009 | rs9662255 | chr1:9441949 |
| rs13238550 | chr7:131059056 | rs4360494 | chr1:38455891 |
| rs1011018 | chr7:139463264 | rs112557609 | chr1:56576924 |
| rs894344 | chr8:135612745 | rs3889199 | chr1:59653742 |
| rs112184198 | chr10:102604514 | rs2289081 | chr2:20881840 |
| rs9549328 | chr13:113636156 | rs11690961 | chr2:46363336 |
| rs9888615 | chr14:53377540 | rs74181299 | chr2:65283972 |
| rs8016306 | chr14:63928546 | rs11689667 | chr2:85491365 |
| rs35199222 | chr15:81013037 | rs1250259 | chr2:216300482 |
| rs11643209 | chr16:75331044 | rs62270945 | chr3:128201889 |
| rs12941318 | chr17:1333598 | rs1566497 | chr4:169717148 |
| rs2467099 | chr17:73949045 | rs17059668 | chr4:174584663 |
| rs6686889 | chr1:25030470 | rs10057188 | chr5:77837789 |
| rs12405515 | chr1:172357441 | rs11154027 | chr6:121781390 |
| rs12408022 | chr1:217718789 | rs36083386 | chr6:152397912 |
| rs10916082 | chr1:227252626 | rs449789 | chr6:159699125 |
| rs2760061 | chr1:228191075 | rs1322639 | chr6:169587103 |
| rs953492 | chr1:243471192 | rs76206723 | chr7:40447971 |
| rs55701159 | chr2:25139596 | rs2978456 | chr8:42324765 |
| rs4952611 | chr2:40567743 | rs4454254 | chr8:141060027 |
| rs76326501 | chr2:43167878 | rs72765298 | chr9:127900996 |
| rs2579519 | chr2:96675166 | rs9337951 | chr10:30317073 |
| rs1438896 | chr2:145646072 | rs10826995 | chr10:32082658 |
| rs79146658 | chr2:179786068 | rs11442819 | chr11:45208141 |
| rs7592578 | chr2:191439591 | rs2289125 | chr11:89224453 |
| rs1063281 | chr2:218668732 | rs8258 | chr11:117283676 |
| rs36022378 | chr3:49913705 | rs139236208 | chr12:94880742 |
| rs743757 | chr3:50476378 | rs9323988 | chr14:98587630 |
| rs9827472 | chr3:56726646 | rs117006983 | chr16:70755610 |
| rs143112823 | chr3:154707967 | rs7500448 | chr16:83045790 |
| rs12374077 | chr3:185317674 | rs7226020 | chr17:6473828 |
| rs66887589 | chr4:120509279 | rs78378222 | chr17:7571752 |
| rs10078021 | chr5:75038431 | rs79089478 | chr17:40317241 |
| rs72812846 | chr5:173377636 | rs62080325 | chr17:42060631 |
| rs13205180 | chr6:51832494 | rs740698 | chr17:60767151 |
| rs147212971 | chr6:166178451 | rs7236548 | chr18:43097750 |
| rs9372498 | chr6:118572486 | rs6081613 | chr20:19465907 |
| rs2978098 | chr8:101676675 | rs12628032 | chr22:19967980 |
| rs62524579 | chr8:144060955 | rs73161324 | chr22:42038786 |
| rs4364717 | chr9:21801530 | rs67330701 | chr11:69079707 |
| rs11030119 | chr11:27728102 | rs7178615 | chr15:66869072 |

Supplementary Table 5: Genetic predictors of lipids included in Mendelian randomization analysis

| rsid | chr:position | rsid | chr:position |
| --- | --- | --- | --- |
| rs10903129 | chr1:25768937 | rs13107325 | chr4:103188709 |
| rs4660293 | chr1:40028180 | rs6450176 | chr5:53298025 |
| rs1998013 | chr1:55958030 | rs9686661 | chr5:55861786 |
| rs10493326 | chr1:62953373 | rs4976033 | chr5:67714246 |
| rs4587594 | chr1:63133930 | rs7703051 | chr5:74625487 |
| rs6603981 | chr1:92993807 | rs4530754 | chr5:122855416 |
| rs12133576 | chr1:93816400 | rs6882076 | chr5:156390297 |
| rs646776 | chr1:109818530 | rs2294261 | chr6:16109163 |
| rs1010167 | chr1:110198727 | rs1800562 | chr6:26093141 |
| rs267733 | chr1:150958836 | rs2247056 | chr6:31265490 |
| rs12145743 | chr1:156700651 | rs205262 | chr6:34563164 |
| rs4650994 | chr1:178515312 | rs998584 | chr6:43757896 |
| rs1689797 | chr1:182150978 | rs17789218 | chr6:100600097 |
| rs2642438 | chr1:220970028 | rs868943 | chr6:116337503 |
| rs903319 | chr1:220985811 | rs9491696 | chr6:127452639 |
| rs4846914 | chr1:230295691 | rs634869 | chr6:139831757 |
| rs6680658 | chr1:230419344 | rs12525163 | chr6:152040291 |
| rs2587534 | chr1:234849339 | rs2297374 | chr6:160575985 |
| rs1367117 | chr2:21263900 | rs1564348 | chr6:160578860 |
| rs515135 | chr2:21286057 | rs702485 | chr7:6449272 |
| rs1260326 | chr2:27730940 | rs17286602 | chr7:16152174 |
| rs3817588 | chr2:27731212 | rs10282707 | chr7:17911038 |
| rs6544713 | chr2:44073881 | rs12670798 | chr7:21607352 |
| rs4148218 | chr2:44099582 | rs4722551 | chr7:25991826 |
| rs2710642 | chr2:63149557 | rs2073547 | chr7:44582331 |
| rs17508045 | chr2:118576719 | rs217386 | chr7:44600695 |
| rs2030746 | chr2:121309488 | rs4917014 | chr7:50305863 |
| rs16831243 | chr2:135762344 | rs17145738 | chr7:72982874 |
| rs7607980 | chr2:165551201 | rs799160 | chr7:73060006 |
| rs355838 | chr2:165619163 | rs38855 | chr7:116358044 |
| rs2287623 | chr2:169830155 | rs3996352 | chr7:130444934 |
| rs1250229 | chr2:216304384 | rs17173637 | chr7:150529449 |
| rs1515110 | chr2:227122216 | rs4240624 | chr8:9184231 |
| rs11563251 | chr2:234679384 | rs9693857 | chr8:9267117 |
| rs9875338 | chr3:12296469 | rs4332136 | chr8:15799853 |
| rs7640978 | chr3:32533010 | rs4921914 | chr8:18272438 |
| rs2290547 | chr3:47061183 | rs12678919 | chr8:19844222 |
| rs2240327 | chr3:50113034 | rs894210 | chr8:19865843 |
| rs13326165 | chr3:52532118 | rs10102164 | chr8:55421614 |
| rs6805251 | chr3:119560606 | rs2326077 | chr8:59385919 |
| rs17345563 | chr3:132209203 | rs2293889 | chr8:116599199 |
| rs687339 | chr3:135932359 | rs2737252 | chr8:116663898 |
| rs1482852 | chr3:156798294 | rs4871137 | chr8:121868551 |
| rs10513688 | chr3:170727218 | rs2980885 | chr8:126474306 |
| rs6831256 | chr4:3473139 | rs2954022 | chr8:126482621 |
| rs10019888 | chr4:26062990 | rs2954022 | chr8:126482621 |
| rs442177 | chr4:88030261 | rs4075205 | chr8:144284709 |
| rs10029254 | chr4:88160140 | rs7832643 | chr8:145022657 |
| rs3822072 | chr4:89741269 | rs3780181 | chr9:2640759 |
| rs2602836 | chr4:100014805 | rs686030 | chr9:15304782 |

Supplementary Table 5: Genetic predictors of lipids (continued)

| rsid | chr:position | rsid | chr:position |
| --- | --- | --- | --- |
| rs7033354 | chr9:16904846 | rs2652834 | chr15:63396867 |
| rs1883025 | chr9:107664301 | rs1035744 | chr15:72566615 |
| rs2472509 | chr9:107684230 | rs3198697 | chr16:15129940 |
| rs8176720 | chr9:136132873 | rs749671 | chr16:31088347 |
| rs579459 | chr9:136154168 | rs9930333 | chr16:53799977 |
| rs1781930 | chr10:5196273 | rs9989419 | chr16:56985139 |
| rs970548 | chr10:46013277 | rs5880 | chr16:57015091 |
| rs7897379 | chr10:65301725 | rs16942887 | chr16:67928042 |
| rs2068888 | chr10:94839642 | rs2288002 | chr16:72057282 |
| rs2255141 | chr10:113933886 | rs2000999 | chr16:72108093 |
| rs2923084 | chr11:10388782 | rs2925979 | chr16:81534790 |
| rs2303975 | chr11:14276999 | rs314253 | chr17:7091650 |
| rs10832962 | chr11:18656271 | rs4791641 | chr17:8161149 |
| rs326214 | chr11:47298360 | rs931992 | chr17:37821435 |
| rs17788930 | chr11:47752775 | rs8077889 | chr17:41878166 |
| rs11246602 | chr11:51512090 | rs7225700 | chr17:45391804 |
| rs12226802 | chr11:55324308 | rs4148005 | chr17:66882466 |
| rs174532 | chr11:61548874 | rs4969178 | chr17:76388202 |
| rs1535 | chr11:61597972 | rs4939883 | chr18:47167214 |
| rs12801636 | chr11:65391317 | rs11660468 | chr18:47209143 |
| rs499974 | chr11:75455021 | rs952044 | chr18:57798110 |
| rs10790162 | chr11:116639104 | rs2278236 | chr19:8431581 |
| rs603446 | chr11:116654435 | rs6511720 | chr19:11202306 |
| rs7117842 | chr11:122534504 | rs688 | chr19:11227602 |
| rs11220462 | chr11:126243952 | rs10401969 | chr19:19407718 |
| rs11045163 | chr12:20463526 | rs731839 | chr19:33899065 |
| rs3741414 | chr12:57844049 | rs1688030 | chr19:35556744 |
| rs10861661 | chr12:107174646 | rs6859 | chr19:45382034 |
| rs2241210 | chr12:109950144 | rs7254892 | chr19:45389596 |
| rs653178 | chr12:112007756 | rs492602 | chr19:49206417 |
| rs6489818 | chr12:112310580 | rs17695224 | chr19:52324216 |
| rs1186380 | chr12:121376416 | rs103294 | chr19:54797848 |
| rs1169288 | chr12:121416650 | rs364585 | chr20:12962718 |
| rs838876 | chr12:125259888 | rs2328223 | chr20:17845921 |
| rs10773105 | chr12:125283766 | rs7264396 | chr20:34154741 |
| rs4942486 | chr13:32953388 | rs6016381 | chr20:39180436 |
| rs1341267 | chr13:95284980 | rs6065311 | chr20:39724338 |
| rs8017377 | chr14:24883887 | rs1800961 | chr20:43042364 |
| rs4983559 | chr14:105277209 | rs4465830 | chr20:44585420 |
| rs2412710 | chr15:42683787 | rs181362 | chr22:21932068 |
| rs492571 | chr15:44211273 | rs5763662 | chr22:30378703 |
| rs1532085 | chr15:58683366 | rs3761445 | chr22:38595411 |
| rs261342 | chr15:58731153 |  |  |

Supplementary Table 6: Genetic variants used in Mendelian randomization analyses for inflammatory biomarkers

* Effect alleles were updated to the forward strand, and hence in some cases differ from that reported by the original publication.

| Biomarker | rsid | Effect allele * | Association with biomarker | Source |
| --- | --- | --- | --- | --- |
| IL-1 | rs6743376 | C | 0.25 SD increase in IL-1Ra | Freitag, 2015[1] |
| IL-1 | rs1542176 | C | 0.18 SD increase in IL-1Ra | Freitag, 2015[1] |
| IL-6 | rs7529229 | T | 0.086 unit increase in log-transformed IL-6 | IL6R MR Consortium, 2012[2] |
| IL-6 | rs4845371 | T | 0.062 unit increase in log-transformed IL-6 | IL6R MR Consortium, 2012[2] |
| IL-6 | rs12740969 | T | 0.078 unit increase in log-transformed IL-6 | IL6R MR Consortium, 2012[2] |
| Fibrinogen | rs7439150 | A | 0.0313 unit increase in log-transformed fibrinogen | de Vries, 2015[3] |
| TNF-alpha | rs1800629 | A | Increased TNF expression | Chu, 2012[4] |
| CRP | rs1205 | C | 0.207 SD increase in log-transformed CRP | Wensley, 2011[5] |
| CRP | rs3093077 | C | 0.169 SD increase in log-transformed CRP | Wensley, 2011[5] |
| CRP | rs1130864 | A | 0.127 SD increase in log-transformed CRP | Wensley, 2011[5] |
| CRP | rs1800947 | C | 0.232 SD increase in log-transformed CRP | Wensley, 2011[5] |
| ICAM-1 | rs1799969 | G | 24.9 ng/mL increase in sICAM-1 | Paré, 2011[6] |
| ICAM-1 | rs5498 | T | 8.0 ng/mL increase in sICAM-1 | Paré, 2011[6] |
| ICAM-1 | rs1801714 | G | 13.8 ng/mL increase in sICAM-1 | Paré, 2011[6] |
| ICAM-1 | rs281437 | C | 1.8 ng/mL increase in sICAM-1 | Paré, 2011[6] |
| ICAM-1 | rs11575074 | A | 7.3 ng/mL increase in sICAM-1 | Paré, 2011[6] |
| P-selectin | rs6136 | A | 6.63 mg/L increase in concentration | Reiner, 2008[7] |

Supplementary Table 7: Mendelian randomization estimates for conventional cardiovascular risk factors on probable lifetime major depression, moderate only

| Risk factor | Method | Odds ratio (95% confidence interval) | p-value |
| --- | --- | --- | --- |
| CHD | IVW | 1.05 (0.98 to 1.13) | 0.18 |
|  | MR-Egger | 1.05 (0.93 to 1.18) | 0.47 |
|  | (intercept) | 1.00 (0.99 to 1.01) | 0.91 |
|  | Weighted median | 1.05 (0.94 to 1.18) | 0.36 |
| BMI | IVW | 1.11 (0.96 to 1.28) | 0.16 |
|  | MR-Egger | 1.29 (0.91 to 1.83) | 0.15 |
|  | (intercept) | 1.00 (0.99 to 1.00) | 0.35 |
|  | Weighted median | 1.21 (0.94 to 1.56) | 0.13 |
| WHR | IVW | 1.02 (0.83 to 1.25) | 0.85 |
|  | MR-Egger | 0.78 (0.34 to 1.80) | 0.56 |
|  | (intercept) | 1.01 (0.99 to 1.03) | 0.52 |
|  | Weighted median | 0.94 (0.71 to 1.24) | 0.63 |
| SBP | IVW | 1.00 (0.96 to 1.05) | 0.85 |
|  | MR-Egger | 0.98 (0.88 to 1.09) | 0.71 |
|  | (intercept) | 1.01 (0.98 to 1.03) | 0.61 |
|  | Weighted median | 0.99 (0.93 to 1.05) | 0.73 |
| DBP | IVW | 1.00 (0.96 to 1.04) | 0.93 |
|  | MR-Egger | 0.95 (0.87 to 1.03) | 0.22 |
|  | (intercept) | 1.01 (1.00 to 1.02) | 0.17 |
|  | Weighted median | 0.99 (0.93 to 1.05) | 0.74 |
| LDL-cholesterol | IVW | 1.01 (0.94 to 1.09) | 0.82 |
|  | MR-Egger | 0.99 (0.88 to 1.11) | 0.84 |
|  | (intercept) | 1.00 (0.99 to 1.01) | 0.66 |
|  | Weighted median | 0.97 (0.87 to 1.09) | 0.65 |
|  | Multivariable | 1.00 (0.93 to 1.08) | 0.97 |
| HDL-cholesterol | IVW | 0.95 (0.88 to 1.03) | 0.23 |
|  | MR-Egger | 0.96 (0.83 to 1.09) | 0.50 |
|  | (intercept) | 1.00 (0.99 to 1.01) | 0.83 |
|  | Weighted median | 0.91 (0.81 to 1.03) | 0.14 |
|  | Multivariable | 1.00 (0.92 to 1.09) | 0.99 |
| Triglycerides | IVW | 1.14 (1.03 to 1.26) | 0.011 |
|  | MR-Egger | 1.10 (0.93 to 1.31) | 0.26 |
|  | (intercept) | 1.00 (0.99 to 1.01) | 0.64 |
|  | Weighted median | 1.12 (0.97 to 1.29) | 0.13 |
|  | Multivariable | 1.15 (1.04 to 1.27) | 0.009 |

Supplementary Table 8: Mendelian randomization estimates for conventional cardiovascular risk factors on probable lifetime major depression, severe only

| Risk factor | Method | Odds ratio (95% confidence interval) | p-value |
| --- | --- | --- | --- |
| CHD | IVW | 1.06 (0.94 to 1.19) | 0.33 |
|  | MR-Egger | 1.07 (0.88 to 1.29) | 0.51 |
|  | (intercept) | 1.00 (0.99 to 1.01) | 0.93 |
|  | Weighted median | 1.03 (0.91 to 1.18) | 0.62 |
| BMI | IVW | 0.90 (0.76 to 1.07) | 0.24 |
|  | MR-Egger | 1.20 (0.80 to 1.82) | 0.38 |
|  | (intercept) | 0.99 (0.98 to 1.00) | 0.13 |
|  | Weighted median | 0.84 (0.65 to 1.07) | 0.16 |
| WHR | IVW | 0.97 (0.78 to 1.21) | 0.79 |
|  | MR-Egger | 1.62 (0.66 to 3.97) | 0.29 |
|  | (intercept) | 0.99 (0.96 to 1.01) | 0.25 |
|  | Weighted median | 1.00 (0.73 to 1.38) | 0.99 |
| SBP | IVW | 0.98 (0.93 to 1.02) | 0.29 |
|  | MR-Egger | 0.94 (0.84 to 1.04) | 0.25 |
|  | (intercept) | 1.01 (0.98 to 1.03) | 0.45 |
|  | Weighted median | 0.97 (0.91 to 1.03) | 0.39 |
| DBP | IVW | 0.99 (0.95 to 1.04) | 0.70 |
|  | MR-Egger | 1.03 (0.95 to 1.13) | 0.47 |
|  | (intercept) | 0.99 (0.98 to 1.01) | 0.29 |
|  | Weighted median | 0.98 (0.92 to 1.05) | 0.58 |
| LDL-cholesterol | IVW | 1.01 (0.91 to 1.12) | 0.87 |
|  | MR-Egger | 1.06 (0.90 to 1.25) | 0.47 |
|  | Weighted median | 1.04 (0.92 to 1.19) | 0.52 |
|  | (intercept) | 1.00 (0.99 to 1.01) | 0.42 |
|  | Multivariable | 0.97 (0.88 to 1.07) | 0.55 |
| HDL-cholesterol | IVW | 0.99 (0.89 to 1.11) | 0.91 |
|  | MR-Egger | 1.01 (0.85 to 1.21) | 0.87 |
|  | Weighted median | 1.08 (0.92 to 1.27) | 0.34 |
|  | (intercept) | 1.00 (0.99 to 1.01) | 0.78 |
|  | Multivariable | 1.06 (0.95 to 1.18) | 0.32 |
| Triglycerides | IVW | 1.23 (1.08 to 1.39) | 0.002 |
|  | MR-Egger | 1.24 (1.00 to 1.54) | 0.047 |
|  | Weighted median | 1.24 (1.05 to 1.46) | 0.011 |
|  | (intercept) | 1.00 (0.99 to 1.01) | 0.88 |
|  | Multivariable | 1.20 (1.04 to 1.37) | 0.010 |

Supplementary Table 9: Mendelian randomization results for inflammatory biomarkers on probable lifetime major depression, moderate only

| Biomarker | Type | OR (95% CI) | p-value |
| --- | --- | --- | --- |
| IL-1 | Per 1 SD increase in IL-1Ra | 1.00 (0.90 to 1.11) | 0.99 |
| IL-6 | Per unit increase in log-transformed IL-6 | 0.74 (0.58 to 0.93) | 0.012 |
| Fibrinogen | Per unit increase in log-transformed fibrinogen | 0.94 (0.32 to 2.80) | 0.91 |
| TNF-alpha | Per TFN expression increasing allele | 0.99 (0.96 to 1.03) | 0.72 |
| CRP | Per 1 SD increase in CRP | 1.11 (0.98 to 1.25) | 0.11 |
| ICAM-1 | Per 1 SD increase in ICAM-1 | 1.02 (0.99 to 1.05) | 0.16 |
| P-selectin | Per P-selectin concentration increasing allele | 0.99 (0.95 to 1.03) | 0.72 |

Supplementary Table 10: Mendelian randomization results for inflammatory biomarkers on probable lifetime major depression, severe only

| Biomarker | Type | OR (95% CI) | p-value |
| --- | --- | --- | --- |
| IL-1 | Per 1 SD increase in IL-1Ra | 1.14 (1.01 to 1.28) | 0.037 |
| IL-6 | Per unit increase in log-transformed IL-6 | 0.76 (0.58 to 1.00) | 0.050 |
| Fibrinogen | Per unit increase in log-transformed fibrinogen | 0.98 (0.27 to 3.49) | 0.97 |
| TNF-alpha | Per TFN expression increasing allele | 1.00 (0.95 to 1.04) | 0.84 |
| CRP | Per 1 SD increase in CRP | 1.26 (1.09 to 1.46) | 0.0014 |
| ICAM-1 | Per 1 SD increase in ICAM-1 | 1.00 (0.97 to 1.04) | 0.82 |
| P-selectin | Per P-selectin concentration increasing allele | 0.95 (0.91 to 1.00) | 0.09 |
